# *Minos* transposon-mediated transgenesis in the sea urchin *Paracentrotus lividus*

**DOI:** 10.64898/2026.06.05.730382

**Authors:** Filomena Caccavale, Giovanni Annona, Pasquale De Luca, Maria I. Arnone

## Abstract

In the multitude of suitable experimental systems used for functional studies in the field of developmental biology, the sea urchin plays a central role due to its amenability to various methods, including both transient and stable transgenesis. Among others, transposable elements represent powerful tools for generating stable transgenic specimens, and *Minos* transposon turned out to be an excellent genetic tool in marine organisms, despite its efficiency being host-dependent. This study provides new evidence for the activity of *Minos* transposable elements and their stable integration into the genome of the Mediterranean sea urchin *Paracentrotus lividus*. Using the *Minos*-based technology coupled with a fully-automated system used for the qPCR screening of the *Minos* transposon integration, we devised a new pipeline for performing transgenesis-based functional studies in *P. lividus*.

## 1. INTRODUCTION

Among deuterostomes, the sea urchin represents a milestone in developmental studies. The possibility to obtain a large number of transparent embryos with synchronous development, together with a detailed embryological atlas and well-established protocols for microinjection of exogenous DNA and RNA into fertilized eggs, make it an invaluable resource for understanding developmental mechanisms [1–4]. Gene function through perturbation assays has had a dramatic impact in scientific research, improving our understanding on the role of a given gene and intricate gene regulatory networks. Several tools have been successfully developed in sea urchins, including both transient and stable manipulation approaches [5–8]. Transgenesis and transient perturbation procedures have been used for decades as main approaches for the investigation of mechanisms occurring during early development. They include, for example, the use of reporter genes for *cis*-regulatory analysis [3], the microinjection of morpholino antisense oligonucleotides (MOs) to block translation or splicing, and the advanced versions, the Vivo-MOs and the caged MOs [6,7,9,10]. Nevertheless, in order to explore processes involved later in development, a fully stable insertion of exogenous DNA (e.g., regulatory genes, enhancers) into the host genome is necessary. Among others, transposable elements represent a powerful tool for generating stable transgenic specimens widely used for functional approaches [11–14]. Transposons are segments of DNA able to move or “transpose” in the genome through the cut-and-paste activity of the transposase protein. The transposase gene, necessary for the excision and insertion, is included in the so called “autonomous” transposon and is flanked by inverted terminal repeats that are the cutting site of the transposase itself. A typical engineered transposon used for transgenesis, is a non-autonomous transposon (it lacks the coding sequence for a functional transposase) termed “donor”. This element carries functional inverted repeats flanking a DNA functional element that should be integrated into the host genome. Usually, the donor is co-injected into the embryo with a source of transposase, as a construct carrying an active transposase gene termed “helper” or a transposase mRNA. In eukaryotes, different transposable elements, for example *P element, Mos1, Minos, Sleeping Beauty* (*SB*), *piggyBac* (*PB*) and *Tol2* transposons, have been extensively used for transgenic applications [13]. *Minos* transposon is a member of the *Tc1*/*mariner* superfamily, originally described in the fruit fly *Drosophila hydei* [15], in which it is very active [16]. Compared to other transposon-based transgenesis approaches, *Minos* insertions in *Drosophila* genes occur with high frequency and preferentially into introns, thus not interfering with normal gene function. Moreover, the integration takes place randomly in the host genome despite a slight TA dinucleotide preference [16,17]. Since its first application, the *Minos* transposon has been found active in several non-host organisms, including marine animals such as the sea squirts *Ciona intestinalis* [18] and *Ciona savignyi* [19], the amphipod *Pharyale hawaiensis* [20] and the sea urchins *Hemicentrotus pulcherrimus* [21] and *Lytechinus pictus* [14]. Experimental results highlighted a substantial variability in *Minos* activity efficiency related to the host, for example a lower excision frequency and *Minos* transposition activity was observed in sea urchins compared to sea squirts [18,19,21]. A similar trend was also reported in species belonging to the same subphylum [18,19]. Here, we tested the activity of *Minos* transposable elements in the sea urchin *P. lividus*. By setting up an efficient and reproducible transgenesis protocol, we obtained a stable *Minos* transposon’s integration into the host genome and confirmed its persistence through larval development. The stability of the integrated exogenous DNA, injected at the zygote stage, was tested through real time PCR, taking advantage of an integrated robotic facility together with automated workflow protocols developed in house. The transgenesis protocol reported here is an efficient and reproducible tool, of high importance for functional studies or for potential drug screen assays in a swathe of marine organisms.

## 2. MATERIALS AND METHODS

### 2.1 Animal care, spawning, and egg fertilization

Adult *P. lividus* were collected in the Gulf of Naples (Italy) and kept in the animal facility at Stazione Zoologica Anton Dohrn (Naples, Italy). Spawning of animals and eggs was performed as previously described [22].

### 2.2 In vitro synthesis of Minos transposase mRNA

As a template for the synthesis of the *Minos* transposase mRNA, the pBlueSKMimRNA plasmid, a gift from Dr. Michalis Averof, was used (Addgene plasmid # 102535; http://n2t.net/addgene:102535; RRID: Addgene_102535) [23]. The plasmid was linearized upon digestion with NotI enzyme (BioLab), and the mRNA was synthesized *in vitro* using the T7 RNA polymerase (HiScribe T7 Quick, BioLab). Later, the Cap structure analog and PolyA tail were added using the Vaccinia Capping System and the *Escherichia coli* Poly(A) Polymerase kits (BioLab), respectively. All the procedures were carried out following the manufacturer’s instructions. Lastly, the mRNA was purified by precipitation with LiCl_2_. Before microinjection, mRNA integrity was checked on a 1% agarose gel and its concentration was estimated by using a NanoDrop 2000c spectrophotometer.

### 2.3 Transgenesis vector

Reporter constructs used in the present study were opposite oriented, inserted in the *Minos* transposon, and included in a unique transgenic vector (Supplementary figure 1A). The first construct was composed by: *i*) a codon-modified NLS-mCherry coding sequence. Nucleotide modifications were made in order to adapt GC content while averting changes at the amino acid sequence; *ii*) the basal promoter and the enhancer of *P. lividus’* histone H1β, based on Lai and collaborators [24–26]; *iii*) a synthetic PolyA signal. The second construct was composed of: *i*) the EGFP coding sequence; *ii*) a 1455 bp sequence upstream to the ATG of the *P. lividus*’ *Opsin4* gene. This genomic region contains a putative enhancer identified as a conserved region between *P. lividus* and *Strongylocentrotus purpuratus* genomes. This element is in proximity of the *Opsin4* ATG and corresponds to the region spanning from 5’-37,543,344 to 3’-37,544,798 on Scaffold_3433 of the *P. lividus* genome (Supplementary figure 1B) *iii*) a SV40 polyA signal. The transgenesis vector pMi{rev[H1β>NLS-mCherry-sPolyA] _Opsin4>EGFP-sv40pA} was synthesized by Twist Bioscience (USA) (Supplementary figure 1B-C).

### 2.4 Microinjection into fertilized eggs

*P. lividus* eggs were prepared for microinjection as previously described by McMahon *et al*., [1]. Eggs were microinjected immediately after fertilization with approximately 8 pL of injection solution containing 5 ng/µL of donor plasmid, 20 ng/µL of DNA carrier (*S. purpuratus* genomic DNA sheared with *HindIII* enzyme), and 300, 600 or 800 ng/µL of *Minos* transposase mRNA. After microinjection, embryos were reared in 0,22 µm filtered sea water (FSW), at 18°C and allowed to grow until the developmental stage of interest.

### 2.5 Excision assay

The excision activity of the *Minos* transposon was tested in *P. lividus* using a protocol modified from Pavlopoulos and Averof [20]. The plasmid pMi(3xP3-EGFP), gifted by Drs. Michalis Averof & Charalambos Savakis (Addgene plasmid # 102540; http://n2t.net/addgene:102540; RRID:Addgene_102540) was used as donor. The mRNA synthesized *in vitro* from the pBlueSKMimRNA plasmid, as described above, was injected as a source of transposase. Two pools of embryos were injected: the first was injected with a solution containing the donor plasmid together with the *Minos* transposase mRNA, while the second pool was injected with the donor plasmid only. Subsequently, once the embryos reached the blastula stage, the total DNA was extracted from both pools (approximately 100 embryos each), using the protocol described in Sasakura et al., [21]. Then an aliquot of 3,5 µL of DNA, was used as template for PCR amplification, with oligonucleotides for T3 (5’-AATTAACCCTCACTAAAGGG-3’) and T7 (5’-TAATACGACTCACTATAGGG-3’) promoters, under the thermal protocol: 95°C for 5 min; 40 cycles at 95°C for 30 sec, 57°C for 30 sec, 72°C for 3 min; a final elongation step of 5 min at 72°C; PCR products were analyzed on a 2% agarose gel.

### 2.6 Automated genomic DNA extraction and real time quantitative PCR preparation

Genomic DNA was extracted from individual sea urchin larvae injected with a solution containing the donor plasmid pMi{rev[H1β>NLS-mCherry-sPolyA] _Opsin4>EGFP-sv40pA} with and without the *Minos* transposase mRNA. DNA extraction was performed in a 96-well plate using the following protocol adapted from D’Agostino et al. [27] and Kroll et al. [28]. All procedures were performed using the high-throughput liquid handling integrated system (TECAN Freedom EVOware), equipped with the MCA96, Multi-Channel Arm 96-tip pipetting head. Each larva was collected in 2 µL of FSW, incubated in 25 mM NaOH + 0,2 mM EDTA (final reaction volume 12 µL) for 15 min at 95°C and cooled at 4°C for 10 min (in a thermal cycler), and then neutralized with 1/10 volume of Tris-HCl 1M pH7. The detailed automated protocol is reported in Supplementary Material 1.

A volume of 1 µl of extracted DNA was used for the real time quantitative PCR (RT-qPCR). PCR reaction composition and cycling conditions were performed following manufacturer’s instructions (PowerTrack SYBR Green Master Mix, Thermo Fisher Scientific). The plate preparation was performed following an automated workflow, adopting a previously developed protocol [29] with a few modifications (Supplementary material 2). The oligonucleotides used to test *Minos* transposon’s integration were: Fw 5’-CAATTCAGCACTCACGCCC-3’; Rev 5’-CAGGGTCAGCTTGCCGTAG-3’. They amplify a region of the transgenesis vector spanning from the putative enhancer of the *Pl-Opsin4* gene to the EGFP’s coding sequence (cds). A serial dilution of the transgenesis vector (10 pg to 10^−4^ pg) and a mix of DNA extracted from injected samples (1:1 to 1:100) were used to construct two separate calibration curves. Since the efficiency of amplification was different between the two calibration curves, we corrected the estimated number of target copies in our samples as described by Gallup [30].

### 2.7 Immunostaining and imaging

Immunostaining was performed as previously described [22,31] with a few modifications. Embryos and larvae were fixed in 4% paraformaldehyde in FSW for 15 min at room temperature (RT), then rinsed in phosphate buffer with 0,1% Tween-20 (PBST). Samples were blocked for 1 hour at RT in 2mg/mL BSA (Sigma-Aldrich) and 4% sheep serum (Sigma-Aldrich) in PBST. Samples were incubated overnight at 4°C with the primary antibody, rat anti-mCherry (Invitrogen) diluted 1:50. After multiple washes in PBST, samples were then incubated for 1 h at RT in the secondary antibody Alexa Fluor® 488 Goat anti-rat IgG H&L (Thermo Fisher Scientific) diluted 1:1000. Afterwards, embryos and larvae were rinsed several times in PBST and finally DAPI (1 μg/mL) was added to label nuclei. Imaging was performed by using a Zeiss LSM 700 confocal microscope and final images were obtained as maximum intensity projection of a Z-stack using FIJI.

## 3. RESULTS

### 3.1 Excision activity of Minos transposon in P. lividus

In order to address the activity of the *Minos* transposable element in *P. lividus*, a specific excision assay was performed by injecting the donor plasmid pMi(3xP3-EGFP) containing a *Minos* transposon of approximately 2000 bp, together with *Minos* transposase mRNA into fertilized eggs (henceforth referred as “+” sample). As a control, we injected the donor plasmid alone (henceforth referred as “-” sample). Embryos were cultured for 24 hours until they reached the late gastrula stage, pooled and processed for DNA extraction.

The efficiency was then examined with PCR by amplifying the transposon region with T3 and T7 oligonucleotides, which are flanking the transposon on the donor plasmid. From our analysis, the amplicon corresponding to the un-excised plasmid was 2242 bp, while the one resulting from the excision event was 221 bp, as expected (Fig. 1). No PCR product corresponding to the un-excision event was detected analyzing the DNA from embryos in which pMi(3xP3-EGFP) was injected (with or without the *Minos* transposase, left panel in Fig. 1), contrary to the un-injected donor plasmid (right panel in Fig. 1).

**Figure 1.**
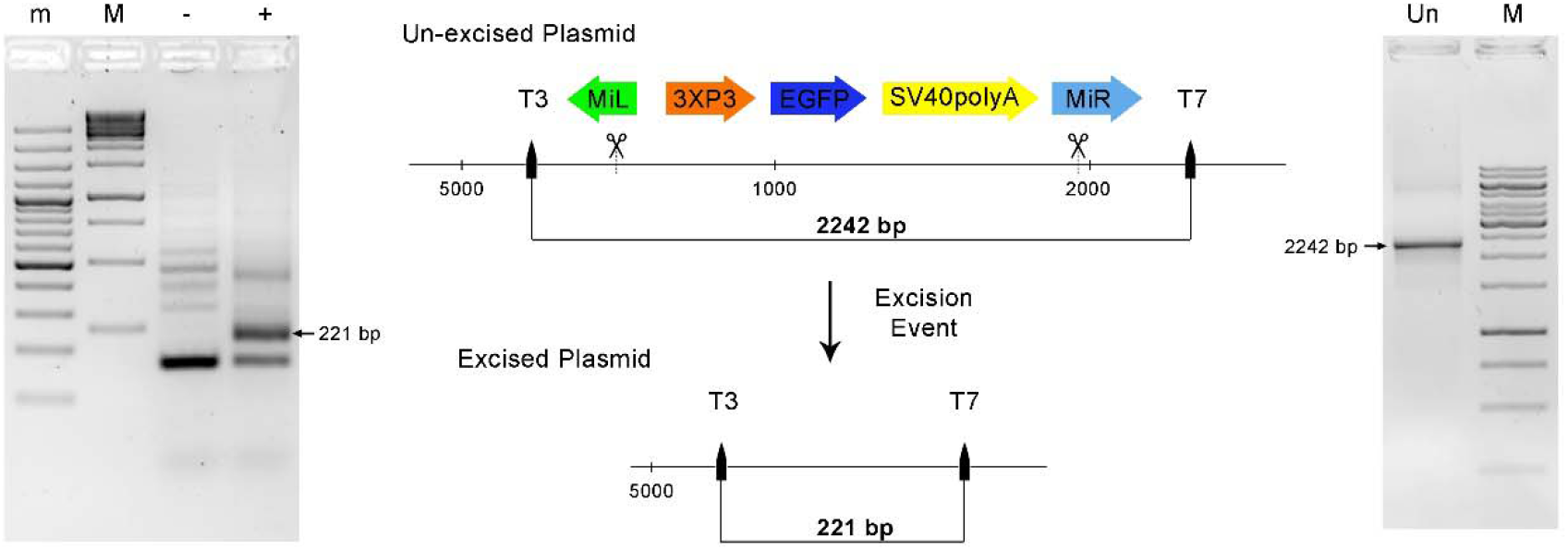
Transposon excision assay. PCR bands derived from: (-) control embryos; (+) embryos in which the donor plasmid was co-injected with the *Minos* transposase; (Un) donor plasmid used as a template. M, GeneRuler 1 kb DNA Ladder; m, GeneRuler 100 bp DNA Ladder.

The PCR analysis derived from both “+” and “-” samples showed bands of various sizes resulting from plasmids shortened by physical damage and/or no specific amplification. Nevertheless, the band corresponding to the excision event was visible only in the “+” sample confirming that *Minos* elements were active in *P. lividus* embryos and that no endogenous *Minos* transposase was present (Fig. 1).

### 3.2 Transposition and genomic integration of Minos transposon into P. lividus embryos

Upon confirmation of the *Minos* transposable element activity in *P. lividus* embryos, we assessed the capability of the system to complete the transposition by re-integrating the excised transposon into the genome. To this aim, the donor plasmid, pMi{rev[H1β>NLS-mCherry-sPolyA] _Opsin4>EGFP-sv40pA}, was co-injected as a circular plasmid, together with 300 ng/µL of *Minos* mRNA. The *Minos* vector was designed with the promoter and enhancer region of the sea urchin late Histone H1β combined with the coding sequence of the fluorescent protein mCherry fused to a nuclear localization signal. This construct was used as a marker for the early identification of transformed specimens. After microinjection, embryos were reared for 24 or 48 hours post fertilization (hpf), corresponding to the late gastrula stage and 2-arm pluteus stage, respectively, and scored for the expression of mCherry. This showed that embryos and larvae co-injected with the *Minos* transposase mRNA and the circular donor plasmid, were positive for nuclear expression of mCherry in a variable number of cells (Fig. 2A-B’). However, we found that mCherry localization was highly variable among embryos of the same batch as well as for different microinjection experiments (Supplementary Figure 2). On the other hand, animals in which the donor plasmid was injected without the source of the *Minos* transposase (control), did not show any mCherry-positive signal (Fig. 2C-D’). The total absence of samples with mCherry-positive cells was also consistently reported in the control group of each microinjection experiment. Moreover, we found that the group of embryos injected with donor plasmid and the *Minos* transposase mRNA, presented variable percentages of mCherry-positive embryos spanning from 23% to 62% in the various microinjection experiments performed (Fig. 2E).

**Figure 2.**
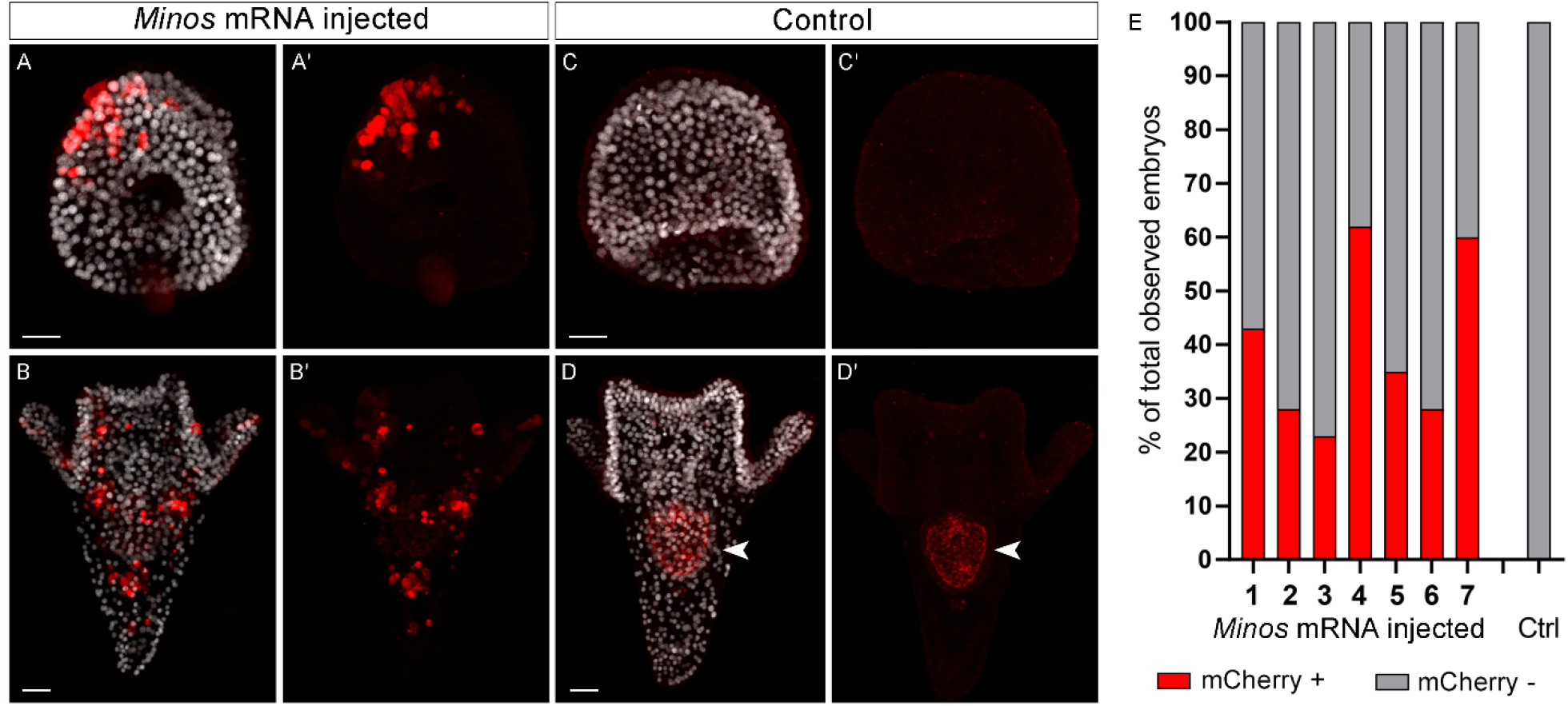
Phenotypic analysis of Minos transposon’s integration into P. lividus’ embryos and larvae. A-B’) Late gastrula and 2-arm pluteus co-injected with donor plasmid pMi{rev[H1β>NLS-mCherry-sPolyA]_Opsin4>EGFP-sv40pA} together with Minos mRNA. C-D’) Control late gastrula and 2-arm pluteus injected with donor plasmid pMi{rev[H1β>NLS-mCherry-sPolyA]_Opsin4>EGFP-sv40pA} without a source of Minos transposase. In A, B, C and D, the nuclei are visualized with DAPI staining and in A’, B’, C’ and D’ the same embryos are depicted without the DAPI channel. The red signal reported in the stomach of the larva in D-D’ (white arrowhead) is due to auto-florescence. E) Percentages of embryos showing the mCherry signal after microinjection of donor plasmid pMi{rev[H1β>NLS-mCherry-sPolyA]_Opsin4>EGFP-sv40pA} alone (Ctrl) or with the source of Minos transposase (Minos mRNA injected). Numbers from 1 to 7 indicate each a different round of microinjection. DAPI: grey; mCherry: red. Scale bars: 20 µm.

In an attempt to understand how the efficiency of transposon’s integration could be increased; we further tested injecting a higher amount of the transposase mRNA (from 300 used before to 600/800 ng). This test led to embryos with lower viability (compared to the ≥ 90% recorded with the 300 ng/µL) and we did not observe any increase in mCherry positive cells (data not shown).

Previous studies have demonstrated that injected DNA that is not integrated into the genome of the zygote is either diluted or degraded during embryonic and larval development [32] while the integrated exogenous DNA remains stable and can be assessed using biomolecular techniques [1,33]. At 1-week post fertilization (4-arm pluteus stage), the target copy numbers present in each larva can be estimated through qPCR, which acts as a confident screening method to assess the stability of the transposon-mediated integration into the genome. As a target, we defined a region of the transgenesis vector spanning from the putative enhancer of the *Pl-Opsin4* gene to the EGFP’s cds. The number of target copies was estimated for single larvae resulting from three different microinjection experiments, identified as microinjection experiments 1, 2 and 3 (Fig. 3). To account for and exclude from our analysis putative residual unintegrated donor plasmid, we also processed in parallel the respective control larvae (not injected) for each experiment. For all the experiments, we estimated that the number of target copies in the control larvae was consistently less than 700, therefore we defined this value (vertical dotted line in Fig. 3) as the baseline threshold to distinguish between larvae with integrated transposon and larvae with no integration. Across the three microinjection experiments, the majority of positive larvae showed an estimated number of target copies between 700 and 5000, with few of them presenting a number of target copies between 7000 and 45000 (Fig. 3). These data corroborated also our qPCR screening results in which we calculated the percentage of positive embryos observed at 24 hpf (Fig. 2E). Furthermore, we observed that the phenotypic analysis of *Minos* transposon’s integration for the same three experiments also was in line with the obtained outcome (Fig. 2E). For microinjection experiment 1, we detected *Minos* transposon’s integration into the genome in 9 out of 25 larvae (36%), compared to 43% of positive embryos observed at 24 hpf (Fig. 2E and 3). For microinjection experiment 2, integration was reported in 21 of 53 larvae (40%), whereas 28% of embryos resulted positive at 24 hpf (Fig. 2E and 3). Regarding the microinjection experiment 3, 18 of the 51 larvae tested (35%) showed *Minos* transposon’s integration, in contrast to 23% of positive embryos observed at 24 hpf (Fig. 2E and 3).

**Figure 3.**
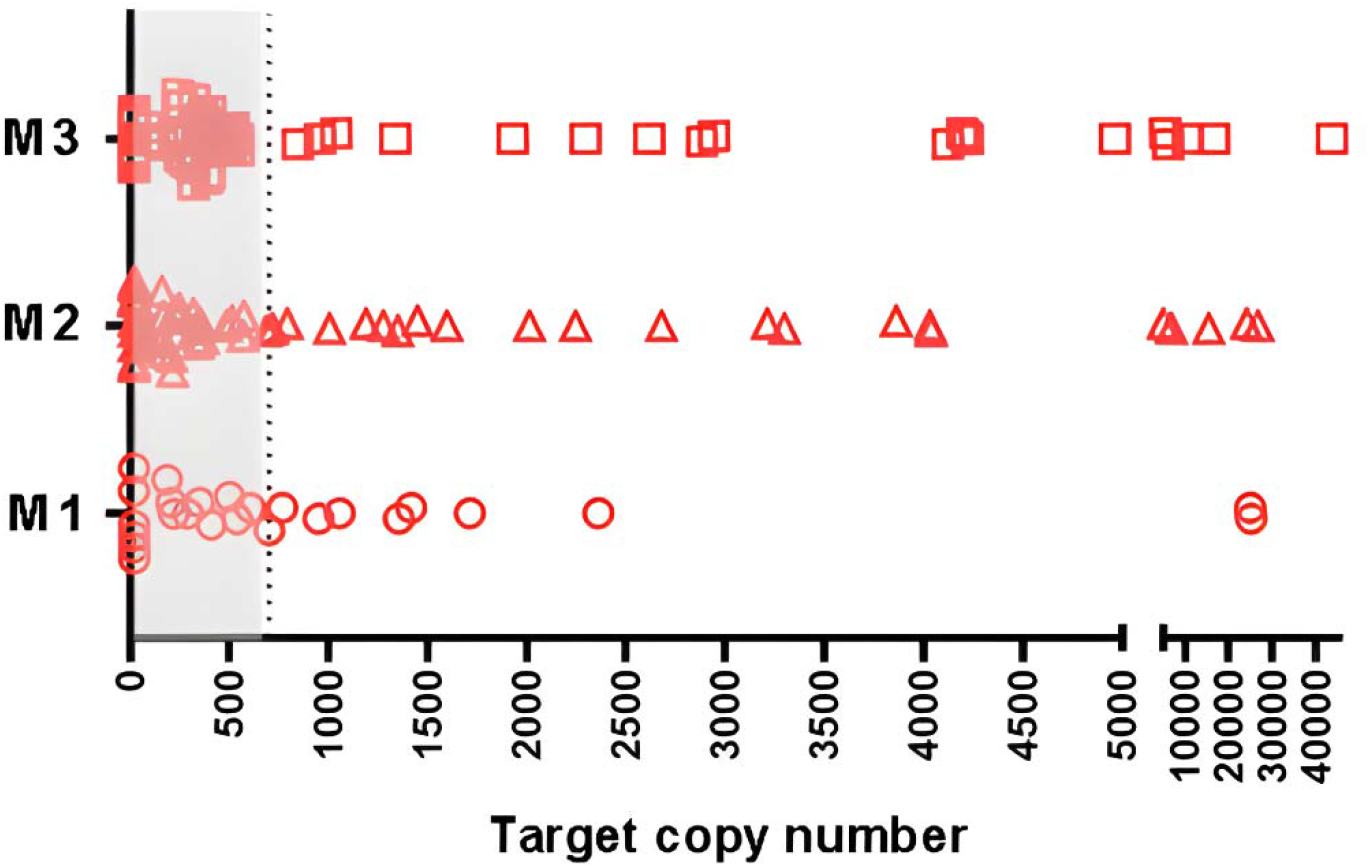
Target copies estimation of *Minos* transposon integrated into the genome of 4-arm plutei 1-week post fertilization. Circles, triangles and squares represent single larvae, obtained from three different batches of embryos and independent injection experiments (M1-M2M3). In each experiment, *Minos* transposase mRNA was co-injected with the donor plasmid pMi{rev[H1β>NLS-mCherry-sPolyA]_Opsin4>EGFP-sv40pA}. The vertical dotted line indicates the baseline threshold between integrated and not integrated *Minos* transposon larvae.

## 4. DISCUSSION

### 4.1 The Minos transposon is active in P. lividus embryos

Transient transgenesis, based on the usage of linearized plasmid constructs, has become a well-established technique for the analysis of *cis*-regulatory elements, across many metazoans including echinoderms. However, most of the methods developed so far are used to understand and map developmental processes taking place during early embryonic development. The aim of the present study was to establish the *Minos* transposon-mediated system as an efficient tool for investigative functional genomic studies in *P. lividus* at a developmental window past embryogenesis, during late larval, pre- and post-metamorphic *P. lividus* developmental stages.

Previous studies have demonstrated that despite a lower efficiency compared to other marine organisms [19,34,34,20], the *Minos* transposon activity not only functions in sea urchins but is also possible to obtain stable transgenic F_1_ lines in the sea urchin species *Lytechinus pictus* [14,21]. Moreover, differently to other transposases such as *Tol2, piggyBAC, Sleeping Beauty* or *Frog Prince, Minos* is currently the only one that is active in sea urchins [14,21].

In detail, Sasakura and collaborators reported that different *Minos* transposase concentrations and culture temperatures did not affect embryogenesis in the sea urchin *H. pulcherrimus*. Furthermore, they observed that higher amounts of *Minos* mRNA (from 200ng/µL to 800ng/µL) increased excision frequency without affecting transposition rate, and that increasing the temperature from 15°C to 20°C did not impact *Minos* transposase activity [21].

In our experiments, we first addressed the activity of *Minos* transposon in *P. lividus* embryos by means of *in vivo* excision assays, followed by testing the ability of the system to produce a stable transposition using a properly designed *Minos* vector, pMi{rev[H1β>NLS-mCherry-sPolyA] _Opsin4>EGFP-sv40pA}. Despite the highly variable and batch-dependent efficiency of the *Minos*-mediated transgenesis, we believe that the results presented in this study, constitute a milestone for the generation of transgenic also in the sea urchin *P. lividus*. Based on all our trials we concluded that the optimal concentration of *Minos* mRNA, as a transposase source, was 300ng/µL. This is due to the fact that at this concentration, we not only obtained transgenic embryos in a range between 23 and 62% of total injected embryos but also all the microinjected specimens were >90% viable. Nonetheless, considering the conclusions reported by Sasakura and collaborators for *H. pulcherrimus*, we also tested 600ng/µL and 800ng/µL of *Minos* transposase (data not shown), without altering the concentration of donor plasmid and carrier DNA used. In such conditions, we not only did not observe any increase in the percentage of transgenic embryos but also the microinjected embryos displayed lower viability. One reason for this could be there is a putative increase in transposition frequency and thus in the integration of the *Minos* transposon in the genome. If this is the case this process could interfere with the normal embryonic development and physiology leading to its precocious death. Taken together, our results suggest that the conditions used for our experiments were sufficient to obtain a stable *Minos* transposition in *P. lividus* embryos.

Moreover, in none of our tests did the transposon integration occur during the early cell divisions, contrary to what happens for other organisms [20], and therefore mosaic transgenic expression in the F_0_ is unavoidable. Notably, a key element in our screening strategy was the inclusion of the promoter of the late Histone H1β fused with the CDS of the mCherry fluorescent protein in our *Minos* vector. This allowed us not only to perform a straightforward screening of the transgenic embryos, but also to disentangle the cells in which, the integration was successful. Of note, this strategy is largely used for screening of transgenic animals obtained with different approaches [20,23,35–39]. At the same time, it is worth mentioning that the expression levels of the fluorescent protein were not detectable throughout larval development, making the visual screening not optimal method. Therefore, as highlighted in our experiments, a PCR-based screening combined with the use of reporter fluorescent protein screening is deemed more suitable to confirm the efficiency and the stability of the integration.

### 4.2 Advantages of using automation in the qPCR screening of Minos transposon’s integration

In order to confirm the integration and the stability of *Minos* transposon into the genome of *P. lividus* embryos we performed a high-throughput qPCR screening on single larvae collected from different batches of embryos and microinjection experiments. The screening was performed at the 4-arm pluteus stage, 1-week post fertilization, in order to minimize the signal due to the un-integrated vector (potential not integrated template). The vector was injected as circular supercoiled DNA, which is known not to integrate into the genome nor to replicate inside the cell [1]. For this reason, in the 1-week-old larvae, we did not expect to retrieve any unintegrated vector that could be, eventually, highly diluted during cell divisions. Such a screening was essential in the initial steps of setting up this protocol that allowed us to confirm the effectiveness of the method. However apart from being an efficient qualitative control it also turned out to be an even more sensitive approach than the early screening through the visualization of the fluorescent markers.

During a PCR-based screening, the contamination of specimens (e.g. with exogenous DNA) is a common and persistent problem that could lead to misinterpretations [40]. To avoid such issues often introduced during highly repetitive steps (such as DNA extraction and 384 well-plate loading), we decided to adopt a fully-automated and time-saving approach. This allowed us to reduce the putative contamination and to process a greater number of samples simultaneously, providing high quality data and reproducibility. Lastly, the data present in this study highlight how high-throughput platforms can be integrated into modernizing and improving laboratory protocols, spanning from the trivial molecular biology methods to high-throughput screening procedures [29,41–44].

## Acknowledgments

We thank Dr. Michalis Averof, for his help in transgenesis vector design and for the gift of the pBlueSKMimRNA plasmid; Dr. Michalis Averof and Dr. Charalambos Savakis for the gift of pMi(3xP3-EGFP) plasmid; Dr. Antonietta Spagnuolo for the gift of the primary antibody anti-mCherry (Invitrogen); Dr. Giovanna Benvenuto, Dr. Enrico D’Aniello, and Dr. Periklis Paganos for critical revision of the manuscript; Paola Cirino and Raffaele Panzuto for animal provisions, and Davide Caramiello for animal maintenance.

