## Supplementary Figure 1 for "*Minos* transposon-mediated transgenesis in the sea urchin *Paracentrotus lividus*"

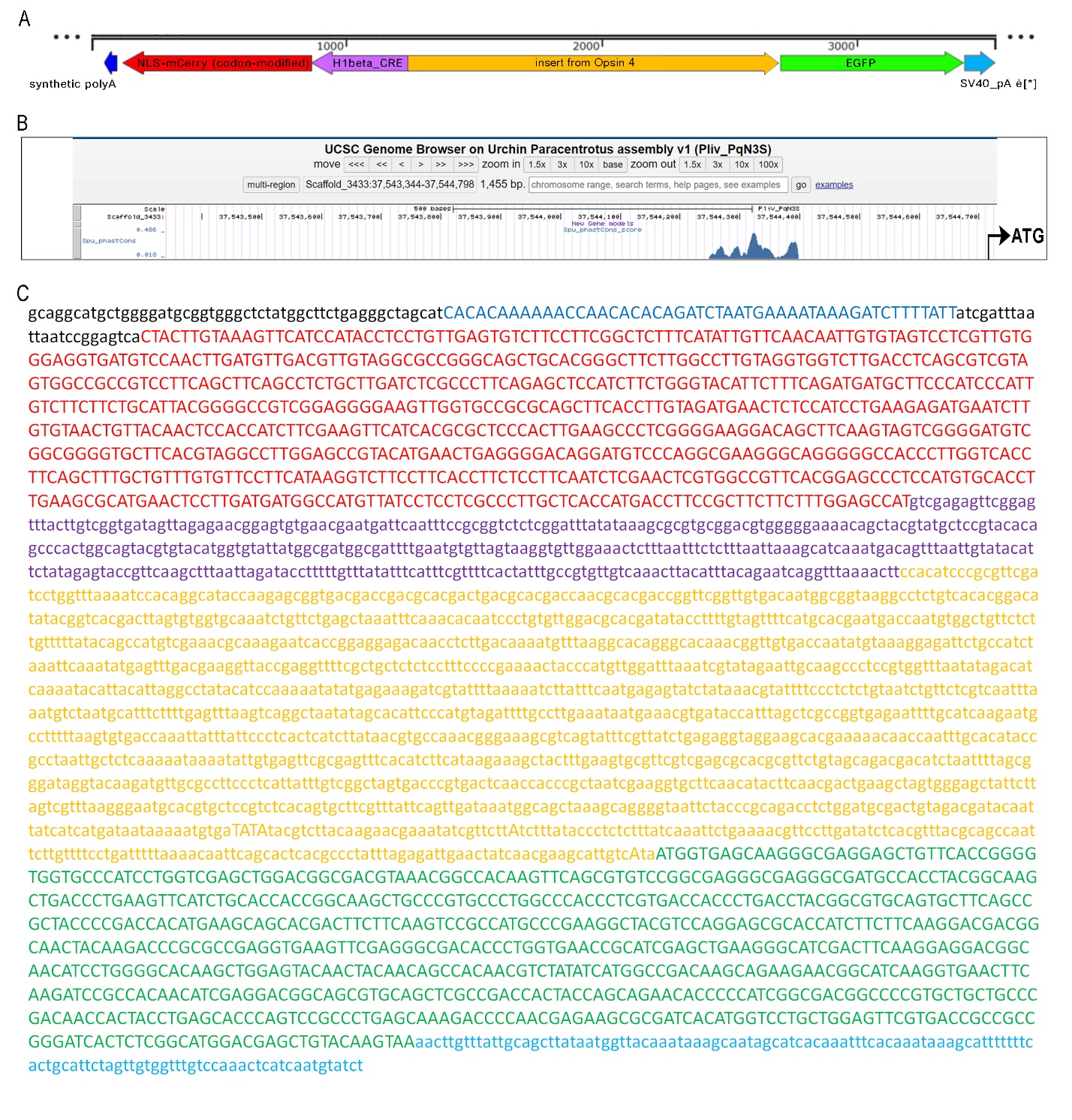

**Supplementary figure 1: Features and sequence of {rev[H1β>NLS-mCherry-sPolyA]_Opsin4>EGFP-sv40pA}, included in the *Minos* donor plasmid has used in the present work.** A) Schematic representation of {rev[H1β>NLS-mCherry-sPolyA]_Opsin4>EGFP-sv40pA}. B) Graphical representation of PhastCons analysis between *Paracentrotus lividus* and *Strongylocentrotus purpuratus* performed on the 1455bp genomic region upstream the ATG of *Opsin4.* Data coming from the UCSC genome browser on urchin *P. lividus* assembly v1 (Pliv_PqN3S), position Scaffold 3433: 37,543,344 - 37,544,798.C) Nucleotide sequence, direction 5’-3’, of {rev[H1β>NLS-mCherry-sPolyA]_Opsin4>EGFP-sv40pA}. Colors correspond with respective features in “A”.
