## Supplementary Figure 2 for "*Minos* transposon-mediated transgenesis in the sea urchin *Paracentrotus lividus*"

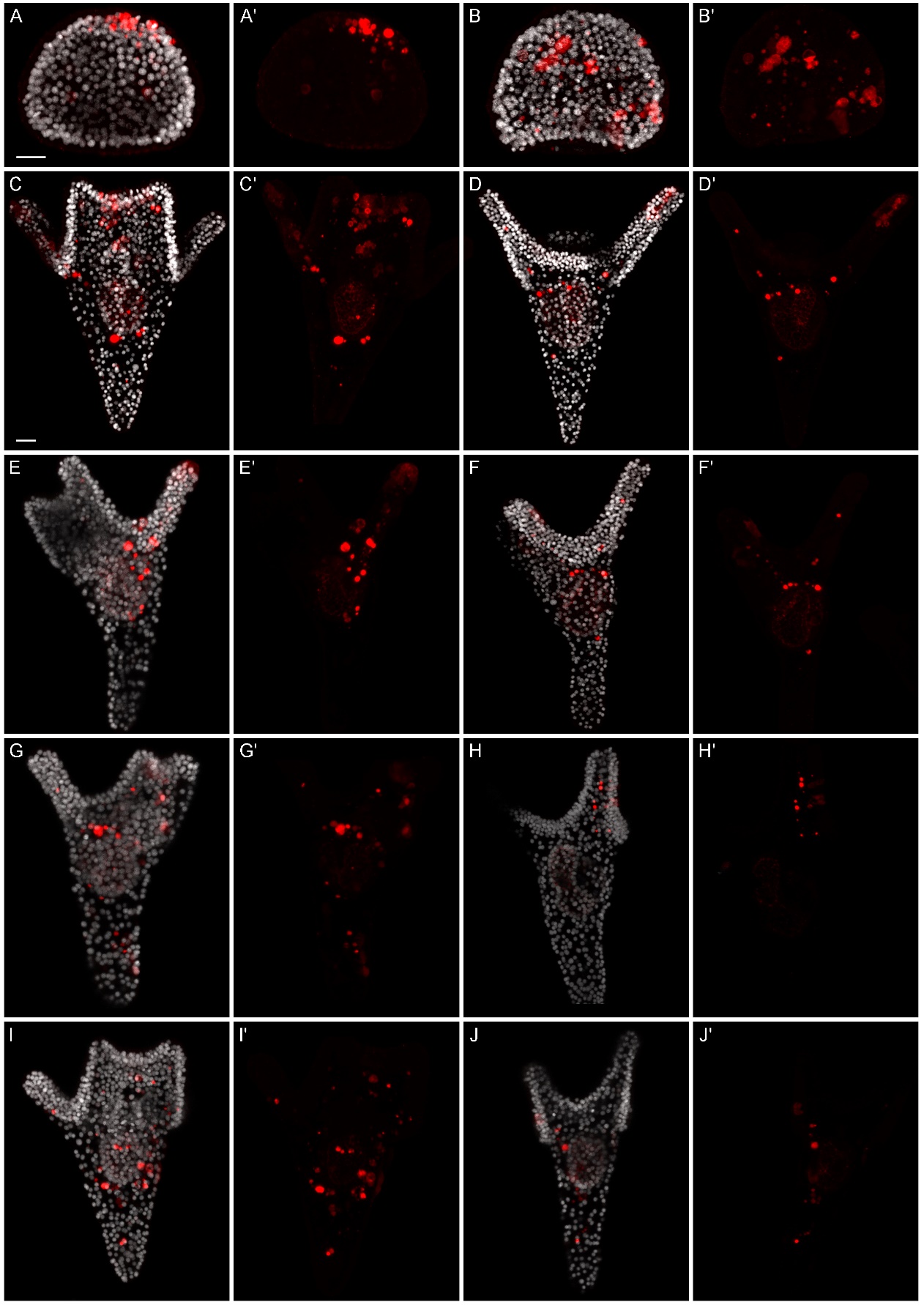

**Supplementary figure 2: Different *P. lividus* specimens showing *Minos* transposon integration by phenotypic analysis.** A-B’) Late gastrula, in A’ and B’ the same embryo of A and B, respectively, showing only mCherry signal. C-J’) 2-arm pluteus, in C’, D’, E’, F’, G’, H’, I’and J’ the same pluteus of C, D, E, F, G, H, I and J, respectively, showing only mCherry signal. Nuclei (DAPI): grey, mCherry: red. Scale bars: 20 µm.
