## Supplementary Material 1 for "*Minos* transposon-mediated transgenesis in the sea urchin *Paracentrotus lividus*"

| No. | Command | Details |
| --- | --- | --- |
| 1 | Wash Tips | 1-8 30 + 40 ml |
| 2 | Get Head Adapter | Grid 62; Site: 2 (Adapter 96 DiTi 4to1 MCA384) |
| 3 | Get DiTis | Grid 50; Site: 1 (DiTi 50ul SBS MCA96)<br>Adapter 96 DiTi 4to1 MCA384 |
| 4 | Aspirate | 10 µl >> Water free dispense Ivan <<<br>"25mM NaOH/0.2mM EDTA" (Col. 1, Rows 1,3,5,7,9,11,13,15) |
| 5 | Dispense | 10 µl Water free dispense Ivan<br>"Larvae in FSW" (Col. 1, Rows 1-8) |
| 6 | Drop DiTis | Grid 50; Site: 1 (DiTi 50ul SBS MCA96)<br>Adapter 96 DiTi 4to1 MCA384 |
| 7 | Drop Head Adapter | Grid 62; Site: 2 (Adapter 96 DiTi 4to1 MCA384) |
| 8 | User Prompt | "Thermocycler step"<br>sound : no |
| 9 | Set DiTi position | DiTi 10ul LiHa<br>Grid : 1, Site : 3, First position in labware : 1 |
| 10 | Begin Loop | 12 times "Tris HCl 1M pH7" |
| 11 | Get DiTis | 1-8 DiTi 10ul LiHa |
| 12 | Aspirate | 1-8 1.2 µl >> Water free dispense Ivan <<<br>"Tris HCl 1M pH7" (Col. 1, Rows 1-8) |
| 13 | Dispense | 1-8 1.2 µl >> Water free dispense Ivan <<<br>"Larvae in FSW" (Col. 1, Rows 1-8) , 1 option |
| 14 | Drop DiTis | 1-8 Washstation 2Grid DiTi Waste |
| 15 | End Loop | "Tris HCl 1M pH7" |

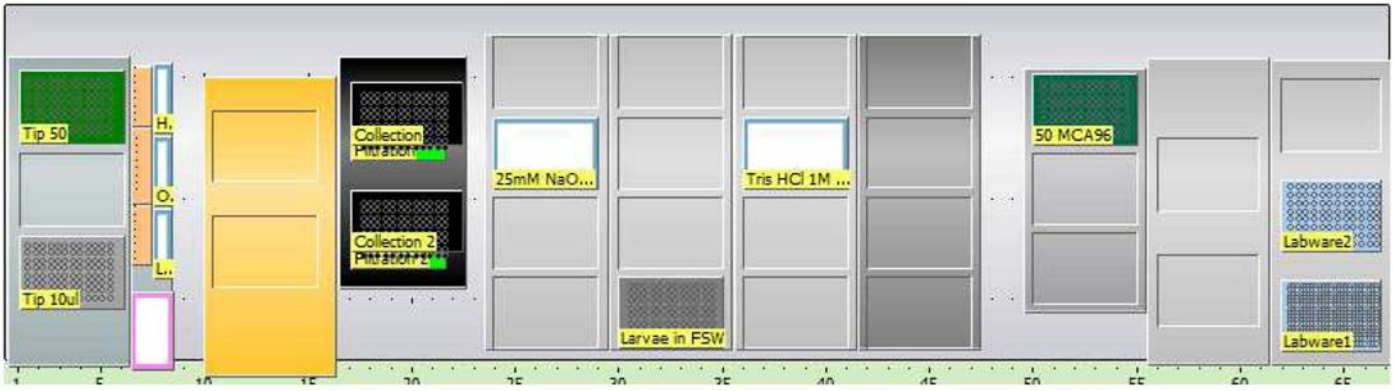
