## Supplementary Material 2 for "*Minos* transposon-mediated transgenesis in the sea urchin *Paracentrotus lividus*"

| No. | Command | Tips | Parameters |
|---|---|---|---|
| 1 | Wash Tips | 1-8 | 30 + 40 ml |
| 2 | Set DiTi position | | DiTi 50ul LiHa<br>Grid : 1, Site : 1, First position in labware : 1 |
| 3 | Get DiTis | 1-8 | DiTi 50ul LiHa |
| 4 | Begin Loop | | 6 times "Mix Epp 1" |
| 5 | Aspirate | 1 | 27 µl >> Water free dispense Ivan << "Master-Mix" (Col. 1, Row 1) |
| 6 | Aspirate | 2 | 27 µl >> Water free dispense Ivan << "Master-Mix" (Col. 1, Row 1) |
| 7 | Aspirate | 3 | 27 µl >> Water free dispense Ivan << "Master-Mix" (Col. 1, Row 1) |
| 8 | Aspirate | 4 | 27 µl >> Water free dispense Ivan << "Master-Mix" (Col. 1, Row 1) |
| 9 | Aspirate | 5 | 27 µl >> Water free dispense Ivan << "Master-Mix" (Col. 1, Row 1) |
| 10 | Aspirate | 6 | 27 µl >> Water free dispense Ivan << "Master-Mix" (Col. 1, Row 1) |
| 11 | Aspirate | 7 | 27 µl >> Water free dispense Ivan << "Master-Mix" (Col. 1, Row 1) |
| 12 | Aspirate | 8 | 27 µl >> Water free dispense Ivan << "Master-Mix" (Col. 1, Row 1) |
| 13 | Dispense | 1-8 | 27 µl Water free dispense Nuova "Mix" (Col. 1, Rows 1-8) , 1 option |
| 14 | End Loop | | "Mix Epp 1" |
| 15 | Drop DiTis | 1-8 | Washstation 2Grid DiTi Waste |
| 16 | Get DiTis | 1-8 | DiTi 50ul LiHa |
| 17 | Begin Loop | | 6 times "Mix Epp 2" |
| 18 | Aspirate | 1 | 27 µl >> Water free dispense Ivan << "Master-Mix" (Col. 2, Row 1) |
| 19 | Aspirate | 2 | 27 µl >> Water free dispense Ivan << "Master-Mix" (Col. 2, Row 1) |
| 20 | Aspirate | 3 | 27 µl >> Water free dispense Ivan << "Master-Mix" (Col. 2, Row 1) |
| 21 | Aspirate | 4 | 27 µl >> Water free dispense Ivan << "Master-Mix" (Col. 2, Row 1) |
| 22 | Aspirate | 5 | 27 µl >> Water free dispense Ivan << "Master-Mix" (Col. 2, Row 1) |
| 23 | Aspirate | 6 | 27 µl >> Water free dispense Ivan << "Master-Mix" (Col. 2, Row 1) |
| 24 | Aspirate | 7 | 27 µl >> Water free dispense Ivan << "Master-Mix" (Col. 2, Row 1) |

| Step | Command | Tips | Parameters |
|---|---|---|---|
| 25 | Aspirate | 8 | 27 µl >> Water free dispense Ivan << "Master-Mix" (Col. 2, Row 1) |
| 26 | Dispense | 1-8 | 27 µl >> Water free dispense Nuova << "Mix" (Col. 7, Rows 1-8) , 1 option |
| 27 | End Loop | | "Mix Epp 2" |
| 28 | Drop DiTis | 1-8 | Washstation 2Grid DiTi Waste |
| 29 | Begin Loop | | 12 times "DNA Larvae" |
| 30 | Get DiTis | 1-8 | DiTi 10ul LiHa |
| 31 | Aspirate | 1-8 | 3 µl >> Water free dispense Nuova << "Larval DNA" (Col. 1, Rows 1-8) , 1 option |
| 32 | Dispense | 1-8 | 3 µl >> Water free dispense Nuova << "Mix" (Col. 1, Rows 1-8) , 1 option |
| 33 | Drop DiTis | 1-8 | Washstation 2Grid DiTi Waste |
| 34 | End Loop | | "DNA Larvae" |
| 35 | Wizard | | ReplicateWizard (...) |
| 36 | Set Variable | | iterationCount = 0 |
| 37 | Begin Loop | | 8 times "LoopSourceCols" |
| 38 | Begin Loop | | 3 times "LoopReplications" |
| 39 | Condition | | iterationCount % 3 <> 0 SkipLabel0_63B6A5EC |
| 40 | Get DiTis | 1-8 | DiTi 50ul Filter LiHa |
| 41 | Comment | | SkipLabel0_63B6A5EC |
| 42 | Aspirate | 1-8 | 10.00 µl >> Water dry contact High density plate << "Mix" (Col. 1, Rows 1-8) , 1 option |
| 43 | Dispense | 1-8 | 10.00 µl Water dry contact High density plate "384 well" (Col. 1, Rows 1-8) , 2 options |
| 44 | Condition | | (iterationCount + 1) % 3 <> 0 SkipLabel1_63B6A5EC |
| 45 | Drop DiTis | 1-8 | Washstation 2Grid DiTi Waste |
| 46 | Comment | | SkipLabel1_63B6A5EC |
| 47 | Set Variable | | iterationCount = iterationCount + 1 |
| 48 | End Loop | | "LoopReplications" |

| No. | Command | Channels | Parameters |
|---|---|---|---|
| 49 | End Loop | | "LoopSourceCols" |
| 50 | Wizard End | | |
| 51 | Wizard | | ReplicateWizard (...) |
| 52 | Set Variable | | iterationCount = 0 |
| 53 | Begin Loop | | 4 times "LoopSourceCols" |
| 54 | Begin Loop | | 3 times "LoopReplications" |
| 55 | Condition | | iterationCount % 3 <> 0<br>SkipLabel0_63B6A637 |
| 56 | Get DiTis | 1-8 | DiTi 50ul Filter LiHa |
| 57 | Comment | | SkipLabel0_63B6A637 |
| 58 | Aspirate | 1-8 | 10.00 µl >> Water dry contact High density plate <<<br>"Mix" (Col. 9, Rows 1-8) , 1 option |
| 59 | Dispense | 1-8 | 10.00 µl Water dry contact High density plate<br>"384 well" (Col. 1, Rows 9-16) , 2 options |
| 60 | Condition | | (iterationCount + 1) % 3 <> 0<br>SkipLabel1_63B6A637 |
| 61 | Drop DiTis | 1-8 | Washstation 2Grid DiTi Waste |
| 62 | Comment | | SkipLabel1_63B6A637 |
| 63 | Set Variable | | iterationCount = iterationCount + 1 |
| 64 | End Loop | | "LoopReplications" |
| 65 | End Loop | | "LoopSourceCols" |
| 66 | Wizard End | | |
| 67 | Group | | Calibration curve |
| 68 | Get DiTis | 1-4 | DiTi 50ul LiHa |
| 69 | Aspirate | 1 | 9 µl Water dry contact High density plate<br>"Master-Mix" (Col. 2, Row 1) |
| 70 | Aspirate | 2 | 9 µl Water dry contact High density plate<br>"Master-Mix" (Col. 2, Row 1) |
| 71 | Aspirate | 3 | 9 µl Water dry contact High density plate<br>"Master-Mix" (Col. 2, Row 1) |
| 72 | Aspirate | 4 | 9 µl Water dry contact High density plate<br>"Master-Mix" (Col. 2, Row 1) |

| | | | |
|---|---|---|---|
| 73 | Dispense | 1 | 9 µl Water dry contact High density plate "384 well" (Col. 21, Row 16) |
| 74 | Dispense | 2 | 9 µl Water dry contact High density plate "384 well" (Col. 22, Row 16) |
| 75 | Dispense | 3 | 9 µl Water dry contact High density plate "384 well" (Col. 23, Row 16) |
| 76 | Dispense | 4 | 9 µl Water dry contact High density plate "384 well" (Col. 24, Row 16) |
| 77 | Drop DiTis | 1-8 | Washstation 2Grid DiTi Waste |
| 78 | Get DiTis | 1-4 | DiTi 50ul LiHa |
| 79 | Aspirate | 1 | 1 µl Water dry contact High density plate "Master-Mix" (Col. 1, Row 4) |
| 80 | Aspirate | 2 | 1 µl Water dry contact High density plate "Master-Mix" (Col. 2, Row 4) |
| 81 | Aspirate | 3 | 1 µl Water dry contact High density plate "Master-Mix" (Col. 3, Row 4) |
| 82 | Aspirate | 4 | 1 µl Water dry contact High density plate "Master-Mix" (Col. 4, Row 4) |
| 83 | Dispense | 1 | 1 µl Water dry contact High density plate "384 well" (Col. 21, Row 16) |
| 84 | Dispense | 2 | 1 µl Water dry contact High density plate "384 well" (Col. 22, Row 16) |
| 85 | Dispense | 3 | 1 µl Water dry contact High density plate "384 well" (Col. 23, Row 16) |
| 86 | Dispense | 4 | 1 µl Water dry contact High density plate "384 well" (Col. 24, Row 16) |
| 87 | Drop DiTis | 1-8 | Washstation 2Grid DiTi Waste |
| 88 | Group End | | Calibration curve |

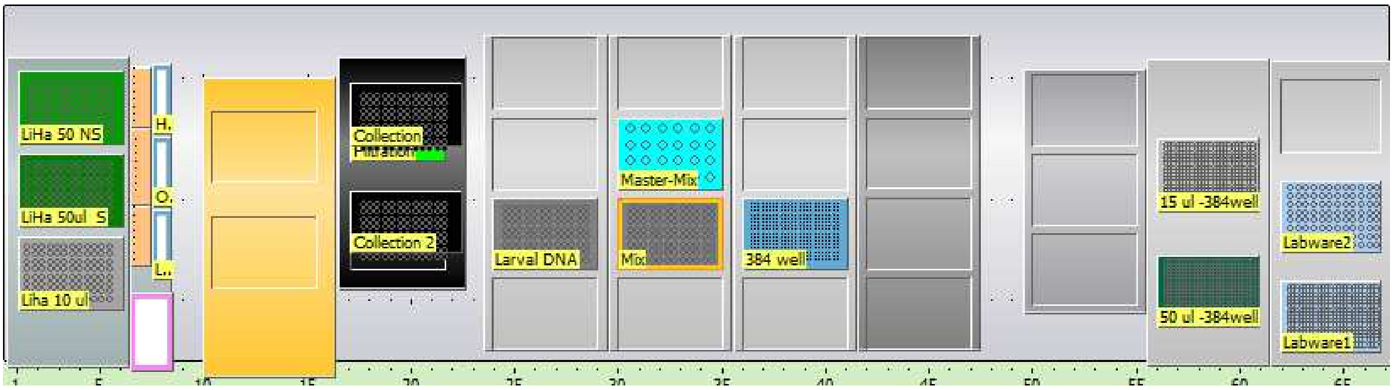
